# SVPopEx: Population-Wide Visualization and Exploration of Structural Variants

**DOI:** 10.64898/2026.08.08.743609

**Authors:** Mari Baker, Kirstin Bett, Ana Vargas, Lingling Jin

## Abstract

Structural variants (SVs) are large-scale genomic variants, which can disrupt important functional and regulatory elements, leading to genomic disorders in humans and playing important roles in domestication, disease resistance, and traits in plants. SVs are generated across populations of individuals and used for association studies, consisting of large datasets with thousands of genomic loci. Visualization of these SVs aids in understanding their genomic distribution, identifying patterns across affected or phenotypic groups, and assessing their proximity to other genomic regions of interest. A variety of tools exist for visualizing SVs, including linear genome browsers and graph-based methods; however, many do not offer intuitive or scalable representations of SVs across large populations. To address this, we present SVPopEx, an interactive tool for population-wide visualization and exploration of SVs. SVPopEx provides a unique and intuitive representation for insertions, deletions, inversions, duplications, and translocations in a linear genome-style browser. Novel features were developed to support comparisons across genomes within user-defined regions, including rendering SVs based on one or more samples and visualizing haplotypes. Use of the tool is demonstrated with SV datasets from *Schistosoma mansoni* and *Lens culinaris*. A task-based evaluation was conducted using SVPopEx and two other linear genome browsers, which demonstrated that SVPopEx excelled in (1) providing a clear representation of the SVs present and (2) supporting comparisons across genomes.

## 1. Introduction

Structural variants are genomic rearrangements greater than 50 base pairs (bp) in length including insertions, deletions, inversions, duplications, and translocations (1). SVs can disrupt functional elements in the genome, leading to effects such as loss of function or altered gene expression, and may influence traits or phenotypes. SVs can also serve as genetic markers in association studies, enabling the identification of relationships between genotypic variation and phenotypic traits.

Variant data is often stored in text-based formats that can contain information for thousands of samples. However, these dense textual representations make it difficult to gain an intuitive understanding of the SVs present within a dataset. Effective visualization of SVs across a population not only provides a clear view of their genomic distribution on a per-sample basis but also reveals population-level patterns and trends. Furthermore, visualizing SVs alongside other genomic features, such as genes and repetitive elements, enhances data interpretation and facilitates the discovery of biologically meaningful insights.

A variety of tools exist for visualizing structural variant calls; however, many fail to provide representations that are both intuitive and scalable for large populations. While many linear genome browsers, such as Integrative Genomics Viewer (IGV) (2) and JBrowse 2 (3), can display population-scale datasets, they often lack meaningful and distinguishable visual representations of different SV types. Graph-based approaches including SequenceTubeMap (4) and svSTM (5) have emerged alongside graph genomes, but they face significant challenges displaying variation, as their node-and-edge structures quickly become cluttered and difficult to interpret. Furthermore, maximally linearized graph visualizations have been criticized for displaying non-meaningful distances between nodes, making it difficult to infer genomic distances from the visualization. Other visualization approaches, such as scatterplots, are effective for displaying copy number variations but are limited in their ability to represent the full spectrum of SV types, restricting their broader applicability.

To address these limitations, we present SVPopEx, a Java-based application for the interactive visualization and exploration of structural variants across populations. SVPopEx provides an intuitive representation of SVs within a scrollable, linear genome-style browser, enabling a natural visualization of genomic coordinates and variant distributions. The tool also facilitates population-level comparisons through user-defined region selection and region-specific analyses. To the best of our knowledge, SVPopEx introduces novel features for comparative rendering of SV calls across multiple samples and haplotype visualization within user-defined genomic regions of interest. Visualization of two datasets with substantially different sizes demonstrates the scalability of SVPopEx and its applicability to diverse organisms and population-scale genomic studies.

## 2. Materials and Methods

SVPopEx is designed to visualize structural variant calls from multiple samples in the context of a reference genome, enabling intuitive exploration and comparison of population-scale genomic variation. Built using the JavaFX 24.0.1 API in Java 24, utilizing only the standard Java and JavaFX libraries, SVPopEx can be easily launched as a standalone desktop application and requires only a multi-sample Variant Call Format (VCF) file as input, providing a simple and streamlined workflow without the need for additional preprocessing or auxiliary files. SVPopEx is implemented using a model–view–controller (MVC) architecture, in which the Model manages data storage and computation, the View provides the graphical user interface, and the Controller coordinates communication between them. Each component is implemented as a separate Java class. The software comprises nine classes: Model, View, Controller, Main, Chromosome, Call, Pair, Sample, and Selection. Their relationships are illustrated in the UML class diagram (Figure 2).

### 2.1 Graphical Interface Design

The graphical interface contains the following components, as illustrated in the schematic representation of the SVPopEx interface, as shown in Figure 1.

**Figure 1.**
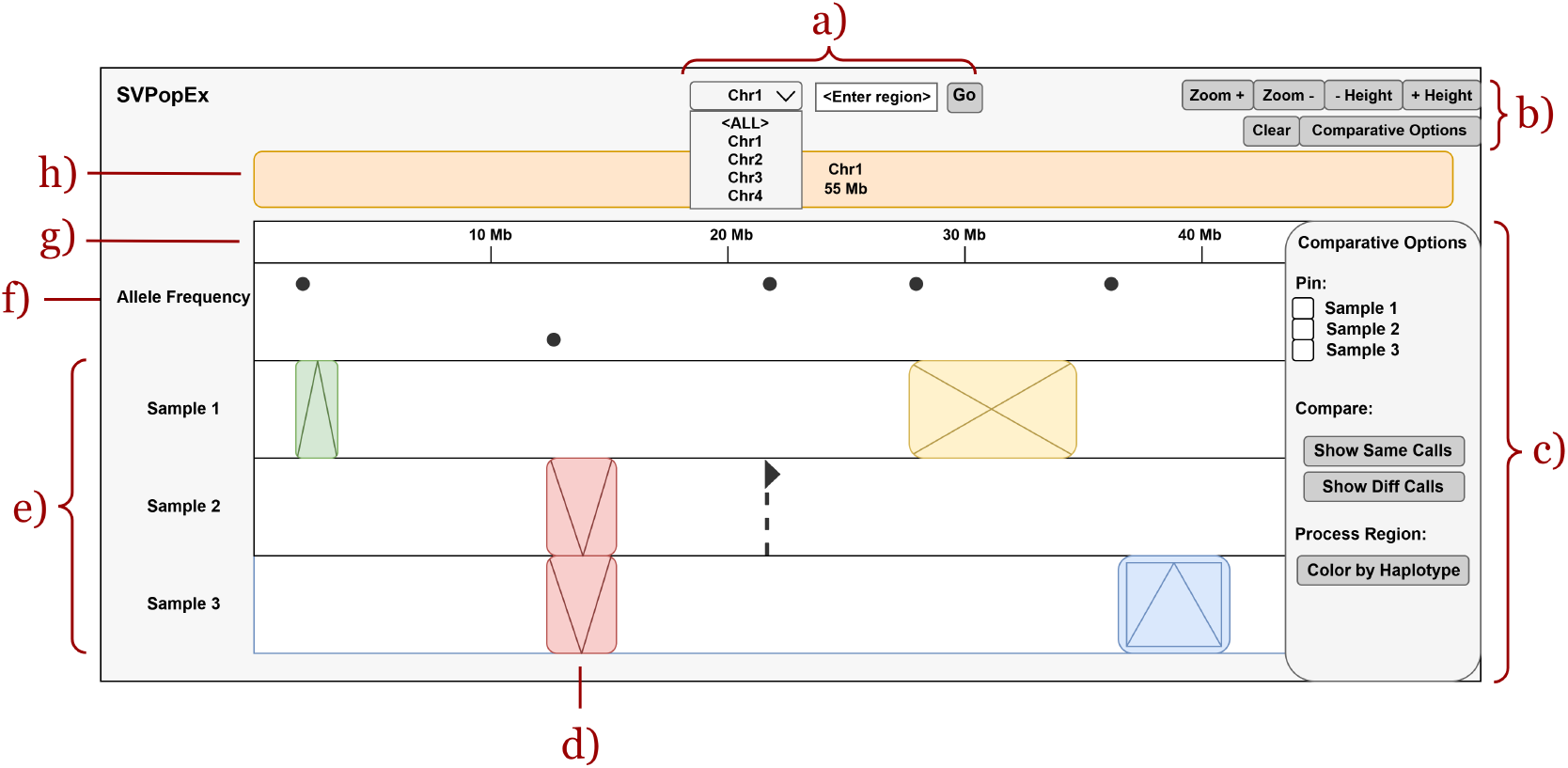
Schematic representation of the SVPopEx interface showing (a) the navigation panel with reference genome selection and custom region input; (b) the variant browser control panel; (c) the side panel for comparative analysis and variant information; (d—g) the variant browser, including (d) structural variant (SV) glyphs, (e) SV tracks, (f) allele frequency track, and (g) genomic coordinates; and (h) the reference genome representation. SV glyphs are encoded as follows: insertions (green rectangles with diverging lines), deletions (red rectangles with converging lines), duplications (blue rectangles with an inner border and diverging lines), inversions (yellow rectangles with crossed diagonal lines), and translocations (black dashed lines with a top triangle).

**Figure 2.**
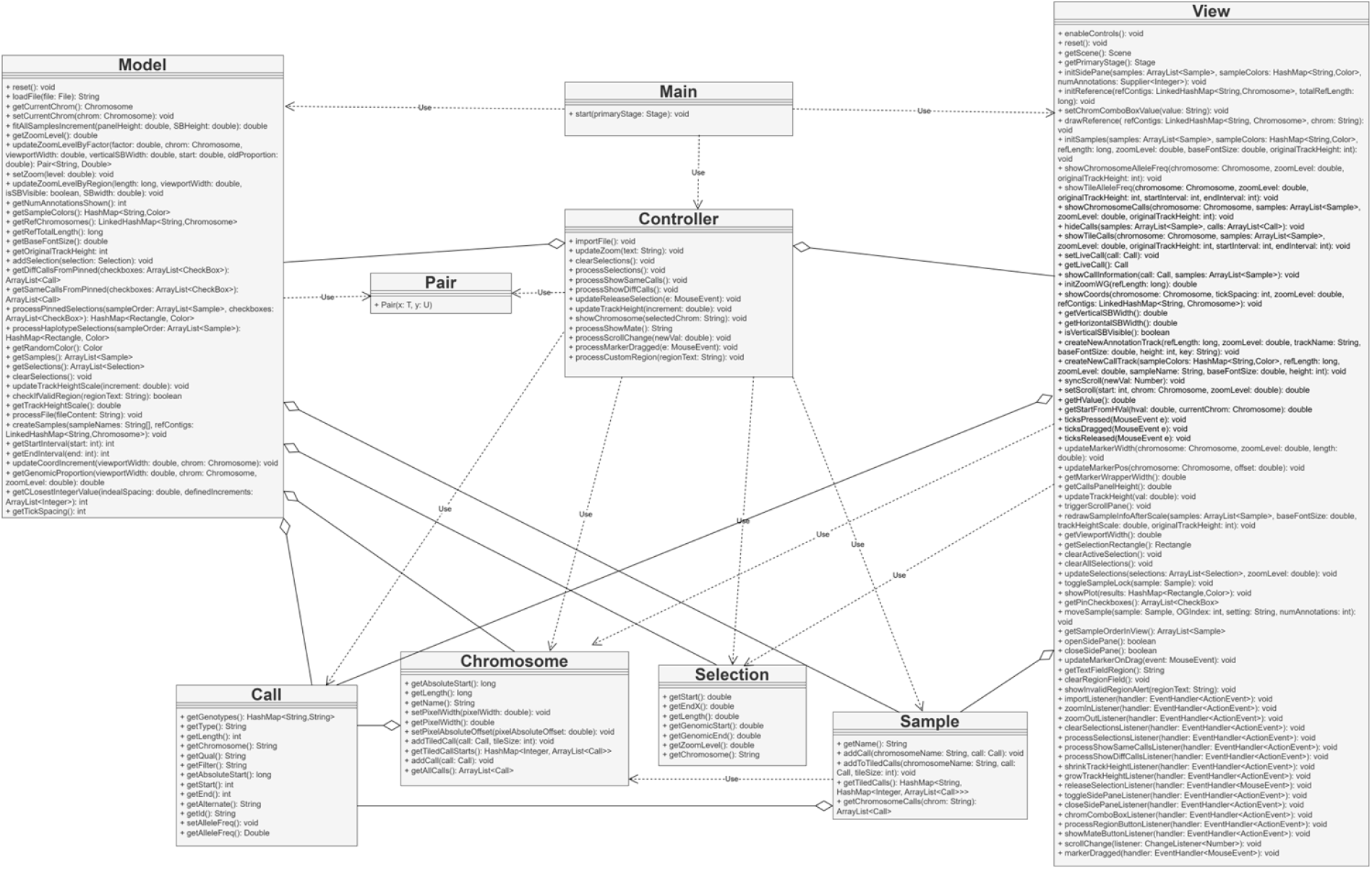
UML diagram of Java classes in SVPopEx shows model-view-controller system design.

#### 2.1.1 SV Glyphs

For each SV in the input VCF file, ‘SVTYPE’ and ‘SVLEN’ are used to generate its graphical representation. SVPopEx supports five SV types: insertion, deletion, duplication, inversion, and translocation.

Each SV type is represented by a distinct glyph that combines unique colors and line patterns for intuitive visual identification. Glyphs are drawn with lengths proportional to ‘SVLEN’ and positioned according to the genomic coordinate specified by ‘POS’.

#### 2.1.2 SV Track

For each sample listed in the VCF header, SVPopEx creates a labeled SV track displaying that sample’s structural variants. An SV glyph is shown at its genomic position if the variant is present in at least one allele. White regions indicate the absence of variant alleles, corresponding to either homozygous reference or missing genotypes. Each sample is also assigned a unique color, which is used by the Color by Haplotype comparative visualization feature.

#### 2.1.3 Allele Frequency Track

The alternate allele frequency (*AF*) for each SV is calculated as

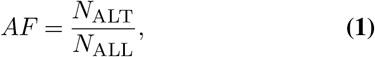

where *N*_ALT_ is the total number of alternate alleles across all samples and *N*_ALL_ is the total number of non-missing alleles. Allele frequencies are computed from the ‘GT’ genotype field of each sample in the input VCF file.

The allele frequency track displays the frequencies of all SVs across the population as a scatter plot, with each point positioned at the SV start coordinate. Frequencies range from 0 at the top of the track to 1 at the bottom, and the background is shaded with a grayscale gradient to facilitate the identification of rare and common variants.

#### 2.1.4 Reference Genome Visualization

SVPopEx uses the contig meta-information lines in the input VCF to obtain the identifiers and lengths of the reference sequences. These sequences, which may represent chromosomes or other genomic assemblies, are displayed in the order specified by the contig entries.

The reference genome is visualized as a bar-like ideogram. In the whole-genome view, each reference sequence is shown as a rounded rectangle with length proportional to its genomic size. An orange overlay, referred to as the view box, indicates the genomic region currently displayed in the variant browser. In the chromosome view, only the selected reference sequence is shown, together with its name and length.

#### 2.1.5 Dynamic Coordinate Display

The genomic coordinate system is displayed beneath the reference genome and aligned with the variant browser tracks, providing positional context for all displayed variants. The coordinate axis updates dynamically during navigation, panning, and zooming to maintain readability and an appropriate density of coordinate labels. In the whole-genome view, it displays chromosome positions across the reference genome, whereas in the chromosome view, it shows genomic coordinates along the selected chromosome. In the chromosome view, users can drag within the coordinate system to select a genomic region for comparative analysis, which is shown as a gray overlay in the variant browser.

#### 2.1.6 Navigation Interface

The navigation interface provides two methods for selecting genomic regions: a reference sequence dropdown menu and a custom region input box. Selecting *<*ALL*>* from the dropdown menu displays the whole-genome view. Custom regions can be specified in the format *<*Chrom:Start-End*>*, where ‘Chrom’ matches the reference sequence identifiers in the input VCF file.

#### 2.1.7 Side Panel Interface

SVPopEx includes a collapsible side panel on the right side of the interface that provides two key functions: comparative analysis options and SV call information. The panel can be expanded or hidden as needed to support interactive exploration of the displayed variants.

#### 2.1.8 Control Panel and Browser Interaction

The control panel provides buttons for interacting with the variant browser and is divided into two groups: general visualization controls (top) and comparative analysis controls (bottom). These controls are disabled in the whole-genome view.

The available controls are:

- Zoom In/Out: Adjusts the horizontal scale of the variant browser by multiplying the current zoom level by 1.5 or 0.5, respectively.
- Track Scaling: Changes the vertical scale of the SV tracks in increments of 0.1.
- Clear: Removes all selections made in the variant browser.
- Comparative Options: Opens the side panel, providing sample-pinning and region comparison tools for down-stream analysis.

### 2.2 Comparative Features

SVPopEx provides comparative analysis tools for exploring structural variants across multiple samples within user-defined regions of interest. Users can select genomic regions, render SVs relative to samples of interest, and visualize haplotypes based on the presence and absence of variants. These features are accessed through the ‘Comparative Options’ button, which opens a dedicated side panel containing sample-pinning and region-based comparison tools.

#### Algorithm 1

Color by Haplotype Plot Algorithm.

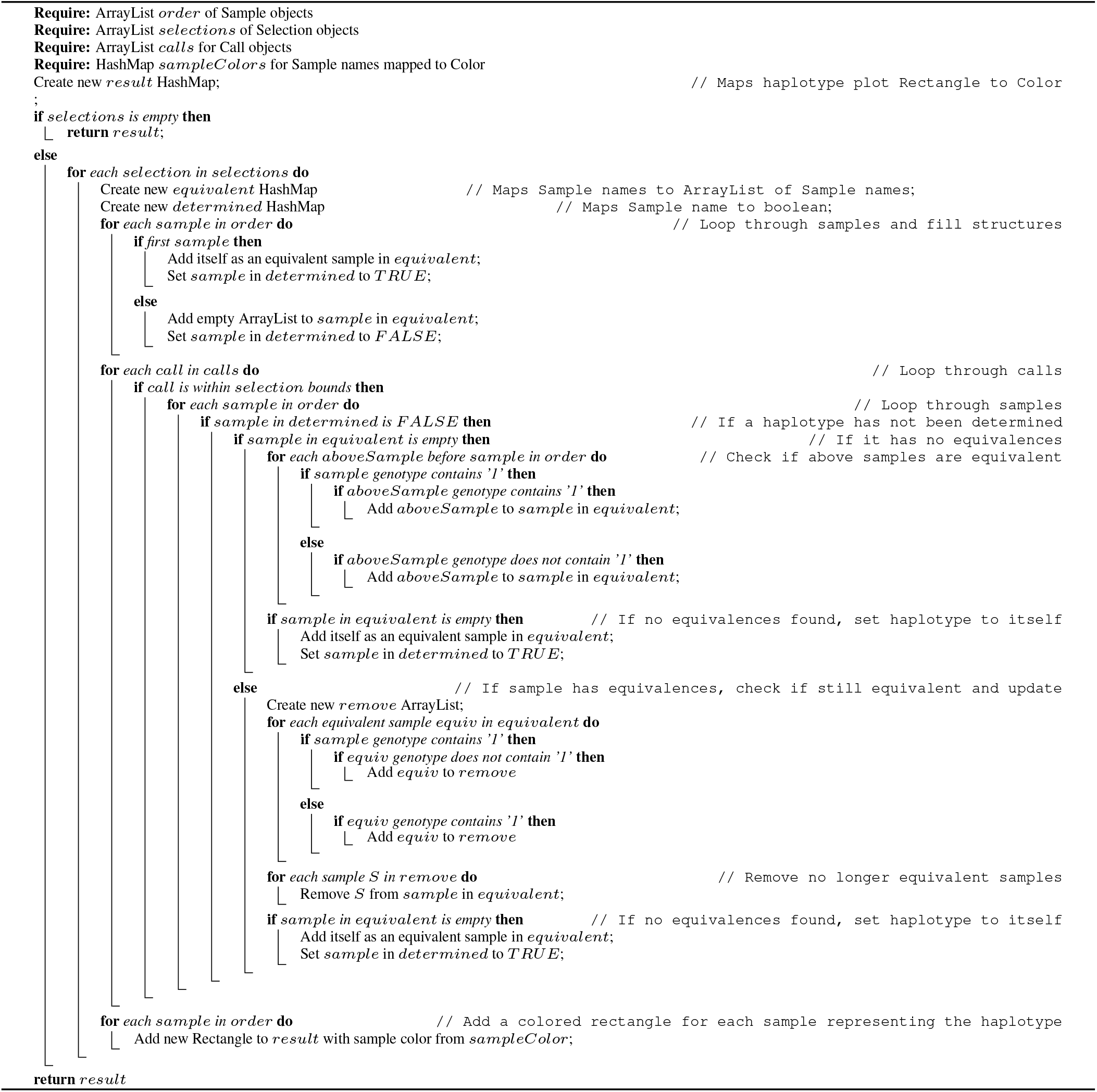

#### 2.2.1 Sample Pinning

Pinning moves selected sample tracks to the top of the variant browser, allowing them to serve as visual references for comparison with other samples. To pin a sample, users open the Comparative Options panel from the control panel and select the corresponding checkbox. Pinned samples are marked with a lock icon and displayed at the top of the browser in the order they are pinned. Unpinning a sample removes the lock and restores it to its original position.

After pinning samples and selecting a genomic region, users can compare structural variants using the ‘Show Same Calls’ and ‘Show Diff Calls’ options in the Comparative Options panel

– Show Same Calls: Highlights SVs present in at least one allele of any pinned sample, while displaying all other SVs with reduced opacity.
– Show Diff Calls: Highlights only SVs that are absent from all pinned samples, rendering all other SVs with reduced opacity.

#### 2.2.2 Region Comparison: Color by Haplotype

The Color by Haplotype feature enables visualization of shared haplotypes among samples within a user-selected genomic region. Haplotypes are based on the presence or absence of SVs for regional SV comparison. After selecting a region of interest, users can activate this function through the Color by Haplotype button in the Comparative Options side panel. Each haplotype is represented by the color of its first matching sample in top-to-bottom display order. The haplotype assignment algorithm is described in Algorithm 1.

### 2.3 Performance Optimization

To improve panning efficiency, SVPopEx adopts a data tiling strategy in which genomic data are partitioned into fixed-size tiles and rendered only when needed. SV calls are assigned to predefined genomic tiles, and only the tiles constituting the current view, together with a small number of neighboring padding tiles, are loaded and displayed during navigation. For each sample, SV calls are indexed by chromosome and tile, enabling rapid retrieval and rendering of the visible region. This approach substantially reduces memory usage by creating and rendering JavaFX nodes only for the required tiles.

## 3. Results

SVPopEx was evaluated using two SV datasets as case studies. Table 1 shows an overview of the two datasets.

**Table 1.** Overview of datasets.

| Organism | *Schistosoma mansoni* | *Lens culinaris* |
| --- | --- | --- |
| Genome Size* | 391 Mb | 3.69 Gb |
| # of Chr | 7 + sex chr | 7 |
| Ploidy | Diploid | Diploid |
| Data Source | NCBI PRJNA1206205 | EVOLVES Project (6) |
| Samples | 5 populations | 23 lines |
| Reads Type | ONT | ONT |
| SV Caller | cuteSV (7) | SVIM (8) |
| # of SVs** | 14,704 | 186,927 |
| SV Types | INS, DEL, DUP, INV, TRA | INS, DEL, INV, BND |
| Load Time | 2.01s | 21.24s |

\*Refers to the assembled reference genome size.
\*\*SV counts reflect the datasets following filtering and exclusions.

### 3.1 Case Study #1

As summarized in Table 1, the dataset comprises SVs from five laboratory populations of *S. mansoni*, generated from publicly available sequencing data by Nair et al. (9) relative to the SM_V10 reference genome (10).

The whole-genome view of the *S. mansoni* populations enables visualization of SV distributions across all chromosomes. Insertions and deletions are substantially more abundant than breakends, inversions, and duplications in the visualization, consistent with the summary statistics in Table 2.

**Table 2.** Distribution of SV types across *S. mansoni* chromosomes.

| Chromosome | DEL | INS | INV | DUP | TRA | Total |
| --- | --- | --- | --- | --- | --- | --- |
| SM_V10_1 | 1,319 | 1,416 | 61 | 26 | 15 | 2,837 |
| SM_V10_2 | 1,337 | 1,317 | 53 | 14 | 4 | 2,725 |
| SM_V10_3 | 1,390 | 1,266 | 30 | 22 | 6 | 2,714 |
| SM_V10_4 | 961 | 1,008 | 30 | 14 | 2 | 2,015 |
| SM_V10_5 | 1,024 | 900 | 34 | 15 | 4 | 1,977 |
| SM_V10_6 | 703 | 710 | 25 | 12 | 4 | 1,454 |
| SM_V10_7 | 465 | 504 | 9 | 4 | 0 | 982 |
| Total | 7,199 | 7,121 | 242 | 107 | 35 | 14,704 |

SV density also varies across the genome, a pattern that is readily apparent in the visualization. The dataset contains abundant SVs that can be effectively compared across populations using SVPopEx.

SVPopEx provides comparative analysis tools for identifying structural variants unique to a population. Figure 4 illustrates SVs unique to the SmOR population on chromosome SM_V10_1. This result is obtained by pinning all populations except SmOR, selecting the genomic region of interest (Figure 3), and applying the Show Diff function to display population-specific SV calls.

**Figure 3.**
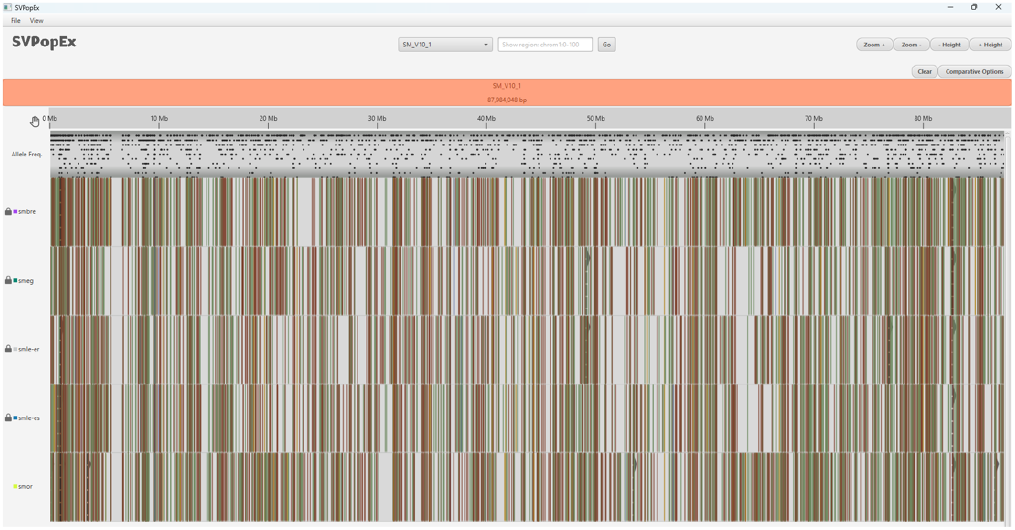
Pinning all *S. mansoni* populations except SmOR and selecting SM_V10_1.

**Figure 4.**
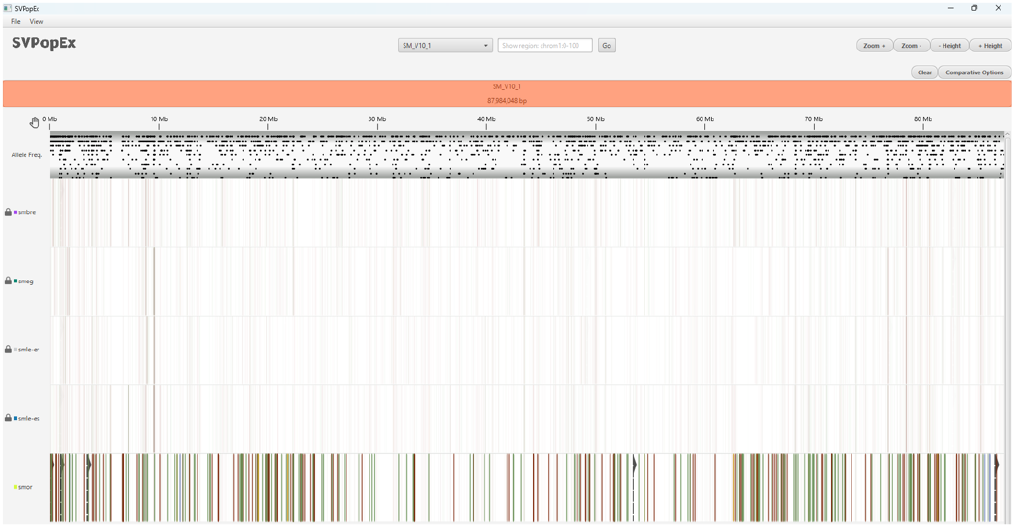
Comparative features in SVPopEx show the unique SVs in the SmOR population on SM_V10_1 by pinning all populations but SmOR and applying the Show Diff function to identify the unique SVs in the SmOR population.

The Color by Haplotype feature summarizes structural variant patterns across genomic regions. Figure 5 shows a 200 kb region containing numerous insertions and deletions, including a particularly dense region between 78.36 Mb and 78.40 Mb.

**Figure 5.**
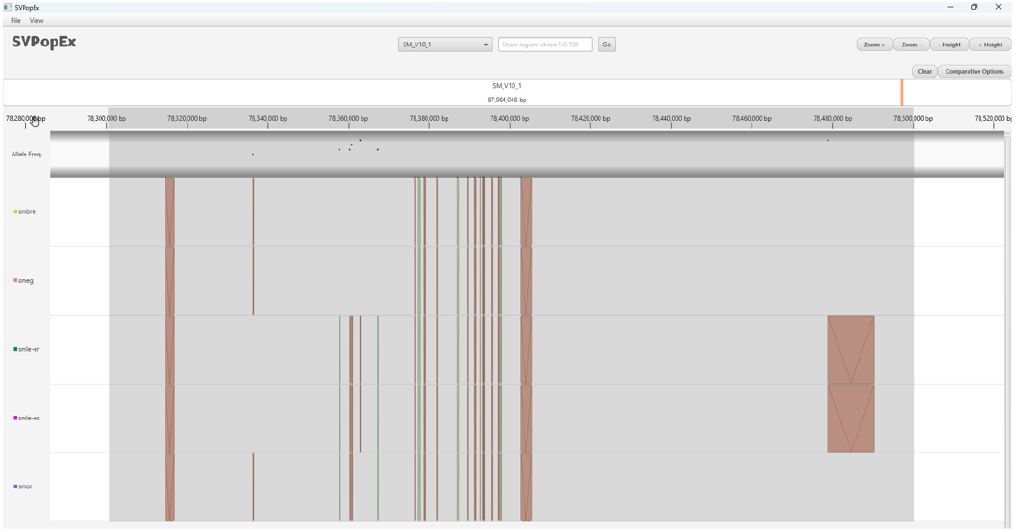
Selection of 78.3–78.5 Mb on SM_V10_1.

The region contains seven genes, four of which encode nucleases. Because the high density of SVs makes visual comparison across populations difficult, the selected region was colored by haplotype (Figure 6) to summarize shared SV patterns. This analysis identified three distinct haplotype groups: (i) SmBRE and SmEG, (ii) SmLE-PZQ-ER and SmLE-PZQ-ES, and (iii) SmOR. The ‘Color by Haplotype’ feature thus provides an intuitive summary of regional SV patterns without requiring manual inspection of individual variants.

**Figure 6.**
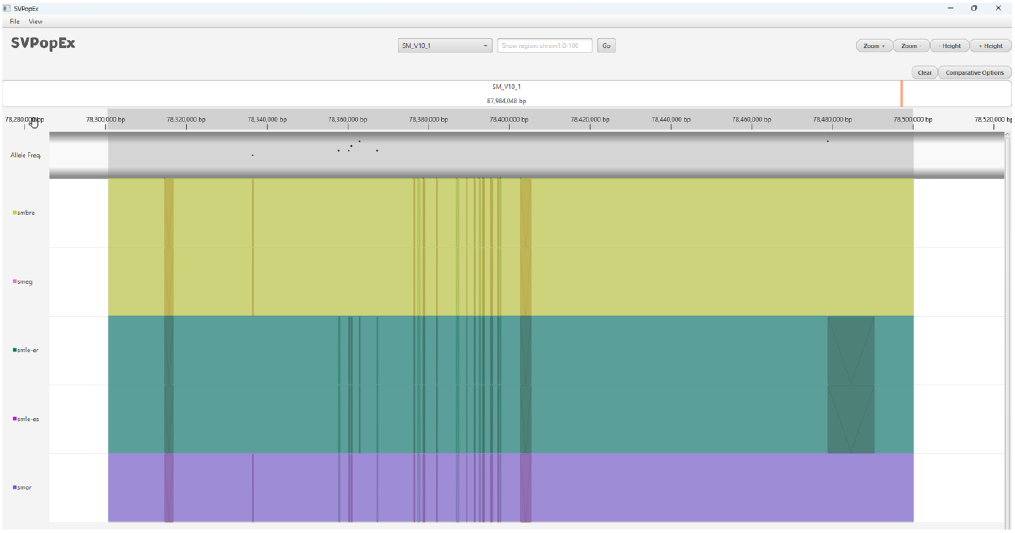
Color by Haplotype identifies three haplotypes within the 78.3–78.5 Mb region of SM_V10_1. Samples are grouped into three haplotypes: (i) SmBRE and SmEG (yellow), (ii) SmLE-PZQ-ER and SmLE-PZQ-ES (blue), and (iii) SmOR (purple).

### 3.2 Case Study #2

Lentil (*Lens culinaris* Medik.) is a legume whose domestication dates back to 11,000 BP in the Fertile Crescent (11, 12). Lentil has seen a global rise in demand and production as a nutritious, fast-cooking, and affordable staple food (13, 14). As shown in Table 1, this dataset consists of SVs identified in 23 lentil lines relative to the Lcu.2RBY reference genome (15).

The whole-genome view of the lentil lines (Figure 7) is dominated by insertions and deletions, consistent with the SV type distribution in Table 3. Reordering the lines using sample pinning reveals shared patterns of variation, including a region with a high deletion density at the end of Lcu.2RBY.Chr2. A higher density of deletions is also observed near the ends of the other chromosomes.

**Table 3.** Distribution of SV types across lentil chromosomes.

| Chromosome | DEL | INS | INV | Total |
| --- | --- | --- | --- | --- |
| Lcu.2RBY.Chr1 | 17,734 | 11,232 | 1 | 28,967 |
| Lcu.2RBY.Chr2 | 22,709 | 12,684 | 3 | 35,396 |
| Lcu.2RBY.Chr3 | 15,660 | 10,186 | 2 | 25,848 |
| Lcu.2RBY.Chr4 | 14,311 | 9,781 | 0 | 24,092 |
| Lcu.2RBY.Chr5 | 13,000 | 9,614 | 3 | 22,617 |
| Lcu.2RBY.Chr6 | 13,820 | 9,296 | 2 | 23,118 |
| Lcu.2RBY.Chr7 | 15,806 | 11,080 | 3 | 26,889 |
| **Total** | **113,040** | **73,873** | **14** | **186,927** |

**Figure 7.**
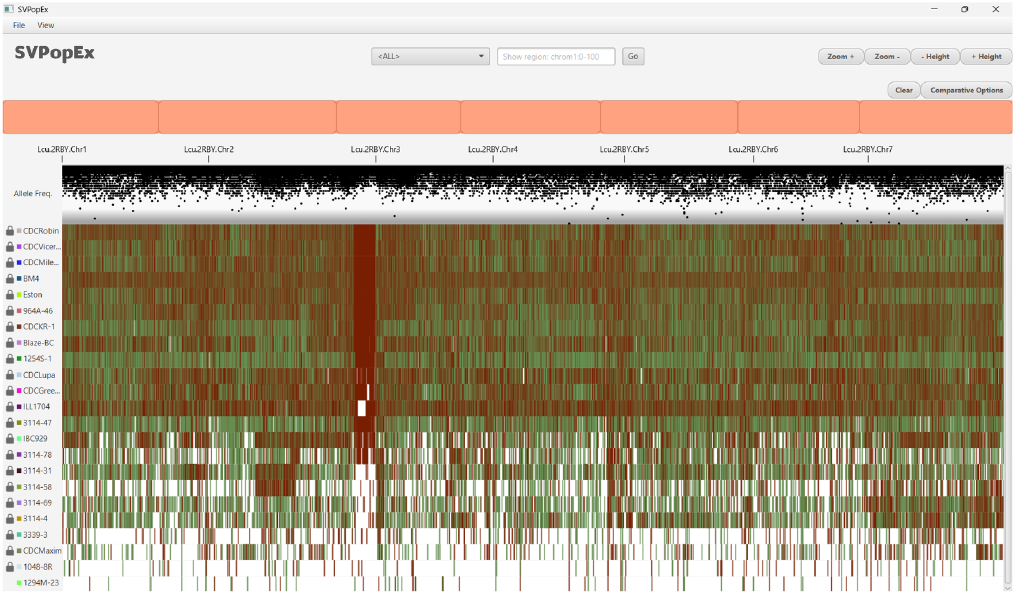
The whole-genome view of the 23 lentil lines. Pinning is applied to reorder lines, revealing variant patterns including a region with a high deletion density at the end of Lcu.2RBY.Chr2.

The allele frequency track in the whole-genome view (Figure 7) reveals that most variants have low allele frequencies (0–0.5), whereas only a small number exhibit frequencies close to 0.9. One such deletion, located at 77.2 Mb on Lcu.2RBY.Chr7 (Figure 8), was further examined by clicking on the variant. The call information panel shows that all lines except CDC Viceroy are homozygous for the deletion allele.

**Figure 8.**
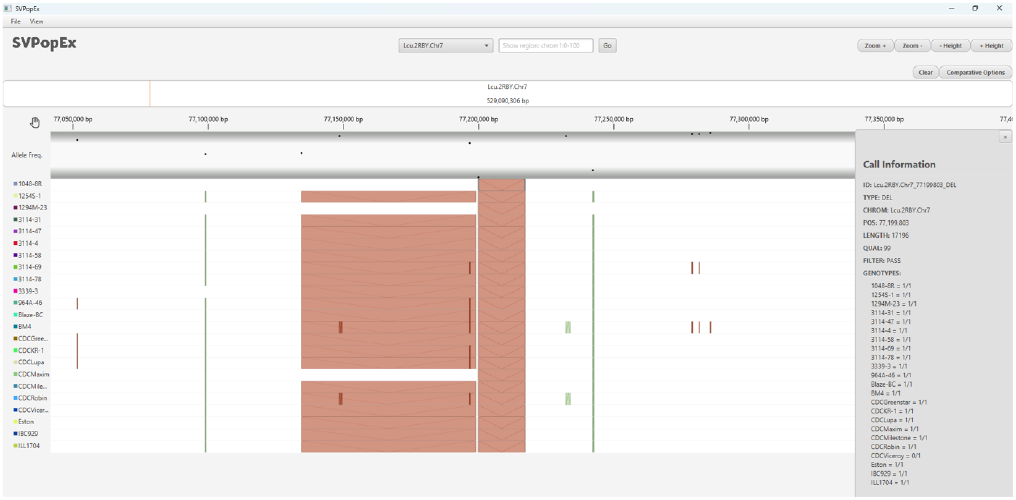
High-frequency deletion variant at 77.2 Mb on Lcu.2RBY.Chr7.

### 3.3 Performance Evaluation

SVPopEx was evaluated on a Windows system with 16 GB of RAM and 4 GB of available Java heap space using the lentil dataset. Heap memory usage was monitored with VisualVM (16) while loading the dataset, zooming to the target region size, and panning for one minute under the test conditions summarized in Table 4. The experiments varied tile size and panning region size to assess their effects on memory consumption and system scalability. The results of the performance evaluation are shown in Figure 9.

**Table 4.** Testing conditions for performance evaluation monitoring heap usage.

| Condition | Tile Size (Mb) | Region Size (Mb) | Padding Tiles | Pan Time (minutes) |
| --- | --- | --- | --- | --- |
| 1 | 5 | 5 | 1 | 1 |
| 2 | 10 | 5 | 1 | 1 |
| 3 | 50 | 5 | 1 | 1 |
| 4 | 5 | 10 | 1 | 1 |
| 5 | 10 | 10 | 1 | 1 |
| 6 | 50 | 10 | 1 | 1 |
| 7 | 5 | 50 | 1 | 1 |
| 8 | 10 | 50 | 1 | 1 |
| 9 | 50 | 50 | 1 | 1 |
| 10 | None | 5 | None | 1 |

**Figure 9.**
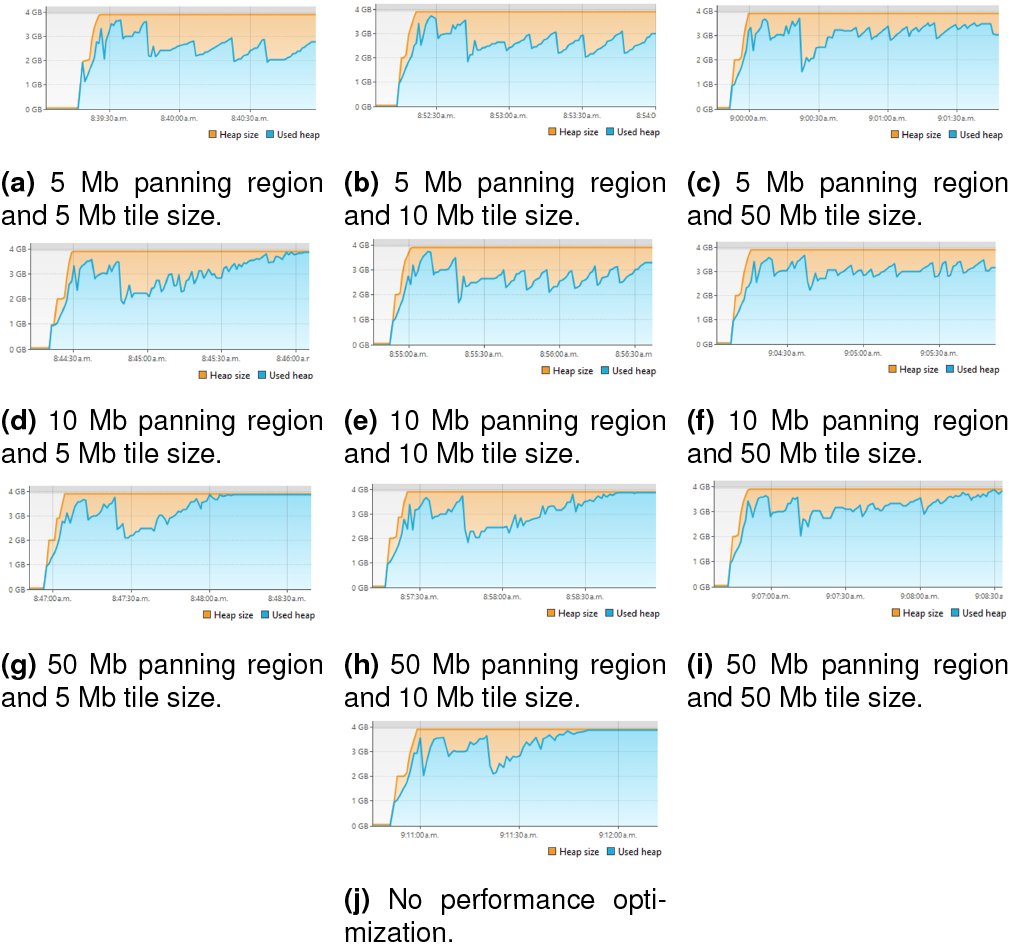
Performance evaluation for lentil dataset showing heap usage for different tile sizes and panning regions. Heap usage was monitored with VisualVM (16) for the test conditions showing (a-b) usage between 2 and 3 GB, (c) usage near 3 GB (d) maximum heap usage and freeze, (e) usage between 2 to 3 GB, (f) usage near 3 GB, and (g-j) maximum heap usage and freeze.

The lentil dataset contains 186,927 SV sites and 615,790 variant alleles across all lines, all of which are stored in memory and rendered as JavaFX nodes. Across all test conditions, loading the VCF file produced an increase in heap usage that peaked between 3 and 4 GB. These results suggest that substantially larger datasets may exceed the available heap space and cause the application to become unresponsive or crash.

Without performance optimization, the application quickly froze, demonstrating the need for efficient memory management when visualizing large datasets. In contrast, the data tiling approach substantially reduced heap usage by rendering only the tiles required for the current view while allowing unused nodes to be garbage collected, thereby improving scalability during interactive navigation.

### 3.4 Task-based Evaluation

A task-based evaluation was conducted to compare the capabilities of SVPopEx with those of the linear genome browsers JBrowse 2 (v4.1.13) (3) and IGV (v2.19.7) (17). The tasks for the evaluation were adapted from the structural variant visualization tool Gremlin (18) for population-wide analysis and are listed in Table 5. The evaluation is based on the tool’s ability to support completion of the tasks through features and interactivity. The task-based evaluation was performed using both the *S. mansoni* and lentil datasets.

**Table 5.** Task-based evaluation comparing IGV, JBrowse 2, and SVPopEx. A dash (**-**) describes no capability, one checkmark (✓) describes severely limited capability or information embedded as text only, two checkmarks (✓✓) describe some capability of the visualization aiding the task but is limited in some capacity, and three checkmarks (✓✓✓) describes full capability.

| Task | IGV | JBrowse 2 | SVPopEx |
| --- | --- | --- | --- |
| T1. Identify the distribution of SVs across the genome. | ✓✓ | ✓✓ | ✓✓✓ |
| T2. Identify the locations of SVs throughout the genome. | ✓ | ✓✓ | ✓✓✓ |
| T3. Identify the variant classification for each SV. | ✓ | ✓✓ | ✓✓✓ |
| T4. Identify the distance and relationship between translocation breakpoints. | ✓ | ✓✓ | ✓ |
| T5. Determine the proximity of SVs to genomic annotations. | ✓✓✓ | ✓✓✓ | - |
| T6. Identify high-confidence SV predictions. | ✓✓✓ | ✓✓✓ | ✓ |
| T7. Determine if multiple genomes exhibit the same SVs. | ✓ | ✓ | ✓✓✓ |
| T8. Identify how and where genomes differ. | ✓ | ✓ | ✓✓✓ |
| T9. Identify the frequency of an SV across genomes. | ✓✓ | ✓ | ✓✓✓ |
| T10. Identify unique SVs for a genome. | ✓ | ✓ | ✓✓✓ |

SVPopEx provides a clearer representation of genome-wide SV distributions than existing tools. IGV summarizes variants as density histograms without preserving sample-level information, whereas JBrowse 2 displays individual variants but suffers from glyph occlusion. In contrast, SVPopEx uses rectangular glyphs with distinct colors and internal designs that remain identifiable even in dense genomic regions, enabling more effective visualization of SV distributions.

SVPopEx also improves the identification of SV types and genomic locations through unique glyph designs for insertions, deletions, duplications, inversions, and translocations. IGV represents SVs using uniform rectangular glyphs that require user interaction to identify the variant class, whereas JBrowse 2 distinguishes only some SV classes and does not scale insertion glyph lengths according to variant size. The explicit glyph design in SVPopEx facilitates rapid visual recognition of variant classes and their genomic positions.

Visualization of translocation relationships remains a challenge across tools. JBrowse 2 supports arc displays that illustrate distances and inter-chromosomal connections between breakpoints, although these views are not available in the multi-sample display and do not distinguish samples individually. IGV and SVPopEx display glyphs for the translocation breakpoints but do not visualize arcs showing the connection between them.

The primary strength of SVPopEx lies in its support for comparative population analysis. Its distinctive glyph design reduces ambiguity between variant types and minimizes occlusion, facilitating comparison of SVs across multiple genomes. Additional features, including haplotype-based coloring, sample pinning, and region-based highlighting of shared or unique variants, further enhance comparative analyses beyond the capabilities of IGV and JBrowse 2.

## 4. Discussion

SVPopEx was developed to provide an intuitive platform for visualizing and exploring structural variants across populations. A key strength of the tool is its glyph-based representation, in which each SV type is distinguished by a unique combination of color, shape, and line patterns. This design effectively conveys SV type, genomic location, size, and distribution while minimizing visual occlusion and distortion. Unlike some linear genome browsers that represent insertions using a fixed glyph length, SVPopEx scales glyphs according to variant size, providing a more faithful representation of SV length. The task-based evaluation further demonstrated that the visualization design facilitates efficient interpretation and comparison of SVs across samples.

Rather than focusing on SV validation through read alignments, SVPopEx is designed for population-scale visualization, comparison, and exploratory analysis of curated SV datasets. As a population-wide visualization tool, SVPopEx complements existing genome browsers such as JBrowse 2, which are better suited for specialized track visualization and variant validation. To improve scalability, SVPopEx employs compact track rendering and a data tiling strategy that reduces the number of graphical nodes displayed simultaneously. These optimizations enable efficient visualization of moderate-sized populations, although the current implementation is limited to approximately 60 samples displayed in the interface before scrolling is required.

Another major contribution of SVPopEx is its support for comparative analysis across user-defined genomic regions. The Color by Haplotype feature summarizes shared combinations of SVs across samples, allowing haplotypes to be visualized according to sample order. Combined with sample pinning and comparative rendering, users can rapidly identify shared and unique SV patterns within regions of interest, which can facilitate the detection of novel and *de novo* variants in datasets.

## 5. Conclusion

Structural variants are a major source of genomic diversity, yet existing visualization tools provide limited support for intuitive population-scale exploration and comparison. This paper presented SVPopEx, a Java-based application that introduces distinct SV glyphs, a linear genome browser with SV and allele frequency tracks, and interactive comparative analysis through sample pinning, region-specific SV comparison, and haplotype visualization. To the best of our knowledge, its region-specific haplotype visualization and sample-based comparative rendering features are novel among current SV visualization tools.

Evaluation on *Schistosoma mansoni* and cultivated lentil datasets demonstrated the utility of SVPopEx for revealing population-level patterns and sample-specific variation, while the allele frequency track enables rapid identification of common and rare variants. Task-based comparisons further showed that SVPopEx provides improved support for identifying, classifying, and comparing structural variants across multiple genomes. Future work will support genome annotation visualization and in-tool SV filtering for enhanced exploration of datasets. SVPopEx complements existing linear genome browsers by providing an intuitive and scalable tool for population-scale SV exploration, making it a valuable resource for comparative genomic research.

## Code Availability

The software and sample dataset is publicly available at:

https://github.com/USask-BINFO/SVPopEx.

## Acknowledgments

This work is funded by Natural Sciences and Engineering Research Council of Canada CREATE grant in Computational Agriculture, Discovery Grant, and Saskatchewan Ministry of Agriculture.

